# “Two-story building” of a ctenophore comb plate provides structural and functional integrity for motility of giant multicilia

**DOI:** 10.1101/2022.03.27.486007

**Authors:** Kei Jokura, Yu Sato, Kogiku Shiba, Kazuo Inaba

## Abstract

Comb plates (CPs) are large compound cilia uniquely seen in a basal metazoan group of ctenophores.^1–3^ Tens of thousands of cilia are bundled in a CP via structures connecting adjacent cilia, called compartmenting lamella (CL), which are the basis for the structural iridescent color and the coordination of ciliary movement of the CP.^4–6^ We previously reported the first component of CL, CTENO64, and found that it was convergently acquired in ctenophores and was essential for determination of ciliary orientation.^3^ However, CTENO64 is localized only in the proximal region of the CL; therefore, the molecular architecture of CL over the entire length has not been elucidated. Here, we identified a second CL component, CTENO189. This ctenophore-specific protein was present in the distal region of the CL, with a localization clearly segregated from CTENO64. Knockdown of the *CTENO189* gene with morpholino antisense oligonucleotides resulted in complete loss of CLs in the distal region, but did not affect either the formation of CP or the orientation of each cilium. However, the hexagonal distribution of cilia was disarranged, and the metachronal coordination of CP along a comb row was lost in the CTENO189 morphants. The morphant CP showed asymmetric ciliary-type movement in normal seawater, and in a high-viscosity solution, it could not maintain the normal waveforms, becoming a symmetric flagellar-type. Our findings demonstrate a “two-story building” of CP, comprising the proximal CL, as the building foundation that rigidly fixes the ciliary orientation. The distal CL would reinforce the elastic connection among cilia to overcome the hydrodynamic drag of giant multiciliary plates.

## RESULTS

### Screening of CL protein candidates

Comb plates (CPs) are iridescent, motile organs that characterize ctenophores. They are giant compound cilia that consist of tens of thousands of cilia with lengths of up to 1 mm.^1,7,8^ All cilia in a comb plate are aligned with the same axonemal orientations. Since the comb plates have fine periodic structures, the light reflected from the comb plates interferes with each other and produces a rainbow structural color.^6^ CP cilia are connected to each other by structures called compartmenting lamellae (CL), which extend from two doublet microtubules (number 3 and 8) of the ciliary axonemes toward the plasma membrane.^1^ The CL is continuous to an extracellular filamentous part, called the interciliary bridge,^4^ where two cilia are physically connected. CLs are thought to be important for the proper orientation and synchronous movement of CP cilia. However, the molecular structures and formation mechanisms of CPs have not been completely elucidated. This is largely because of the lack of information on the CL components.

We reported the first CL protein, CTENO64, in our previous paper, but the protein showed limited localization to only the proximal region of the CP.^3^ To clarify the whole picture of the molecular composition of CL, we searched for other candidates of CL proteins using two criteria (Figure S1A). First, whole CP proteins from the ctenophore *Bolinopsis mikado* were separated by SDS-PAGE, and the gel was cut into 10 slices. Proteins in each slice were analyzed using nanoLC-MS/MS, and 2,052 proteins were identified. Since CL represents a large electron-dense structure, we hypothesized that CL proteins would be present in relatively high amounts. Therefore, the CL proteins of interest were narrowed down to 57 proteins using the exponentially modified protein abundance index, emPAI,^9^ of more than 50. Second, we selected CP cell-specific gene clusters C48 and C50 using published single-cell transcriptome data of *Mnemiopsis leidyi*, a closely related species of *B. mikado*.^10^ We confirmed that CTENO64 was detected in both C48 and C50 cell type clusters. A total of 205 proteins were identified in both clusters (Figure S1A). Finally, we selected 21 proteins based on both criteria (Table S1). A BLASTP search of these 21 CL candidates against known proteins revealed that the protein c56364_g1_i1 was a novel protein with no significant homology to known proteins (Table S2). Local BLAST searches for this protein against published transcriptome data of 11 ctenophore species showed that its homologs were present in all other ctenophore species (Table S3). Therefore, we focused on this protein in this study.

### CTENO189 is a novel CL protein localized in the comb plate distal region

The c56364_g1_i1 protein was composed of 1582 amino acids with a metalloprotease domain (MP) at the amino acid position 223-409 and two death domains (DD) at 1374–1457 and 1493–1570 (Figure S1B). Although a portion of the metalloprotease domain showed homology with those in zinc-dependent metalloproteases, such as astacin and a bacterial and fungal matrix metalloproteinase, the overall sequence of c56364_g1_i1 was not homologous to any known proteins.

To investigate whether CTENO189 is a structural component of CL, we prepared a polyclonal antibody against the recombinant c56364_g1_i1 protein. The antibody specifically recognized a 189-kDa protein in whole comb plate proteins and slightly detected some lower molecular mass proteins, most likely the degradation products of the 189-kDa protein, as the amount changed in different comb plate preparations (Figure S1C). Considering that this 189-kDa protein is also uniquely present in ctenophore species, we named this protein CTENO189.

Immunoelectron microscopy of CTENO189 showed clear immunogold localization in the CL (Figure 1A). Fluorescent double-staining with acetylated α-tubulin showed that CTENO189 was localized in the distal region of the CP both in an adult and a larva, but was not found in other cilia, including those in the pharynx, an apical organ, or ciliated grooves (Figure 1B). We previously identified CTENO64 as a CL protein localized in the proximal region of a CP.^3^ Double-staining with CTENO64 revealed that CTENO189 and CTENO64 were localized to distal and proximal regions of the entire adult CP, respectively, with clear boundaries (Figure 1C). For a more detailed localization analysis, the cilia of CPs were mechanically disintegrated, double-stained with antibodies against both proteins, and observed using a super-resolution lattice-structured illumination microscope (SIM). Double-staining of each CL protein with acetylated α-tubulin showed that both CL proteins were localized along the two lateral rows of an axoneme (Figure 1D). Intriguingly, we further confirmed a clear boundary between CTENO189 and CTENO64 at the level of one axoneme, indicating that the two proteins were separately localized without any overlap along the row of a CL (Figures 1E and S2).

**Figure 1.**
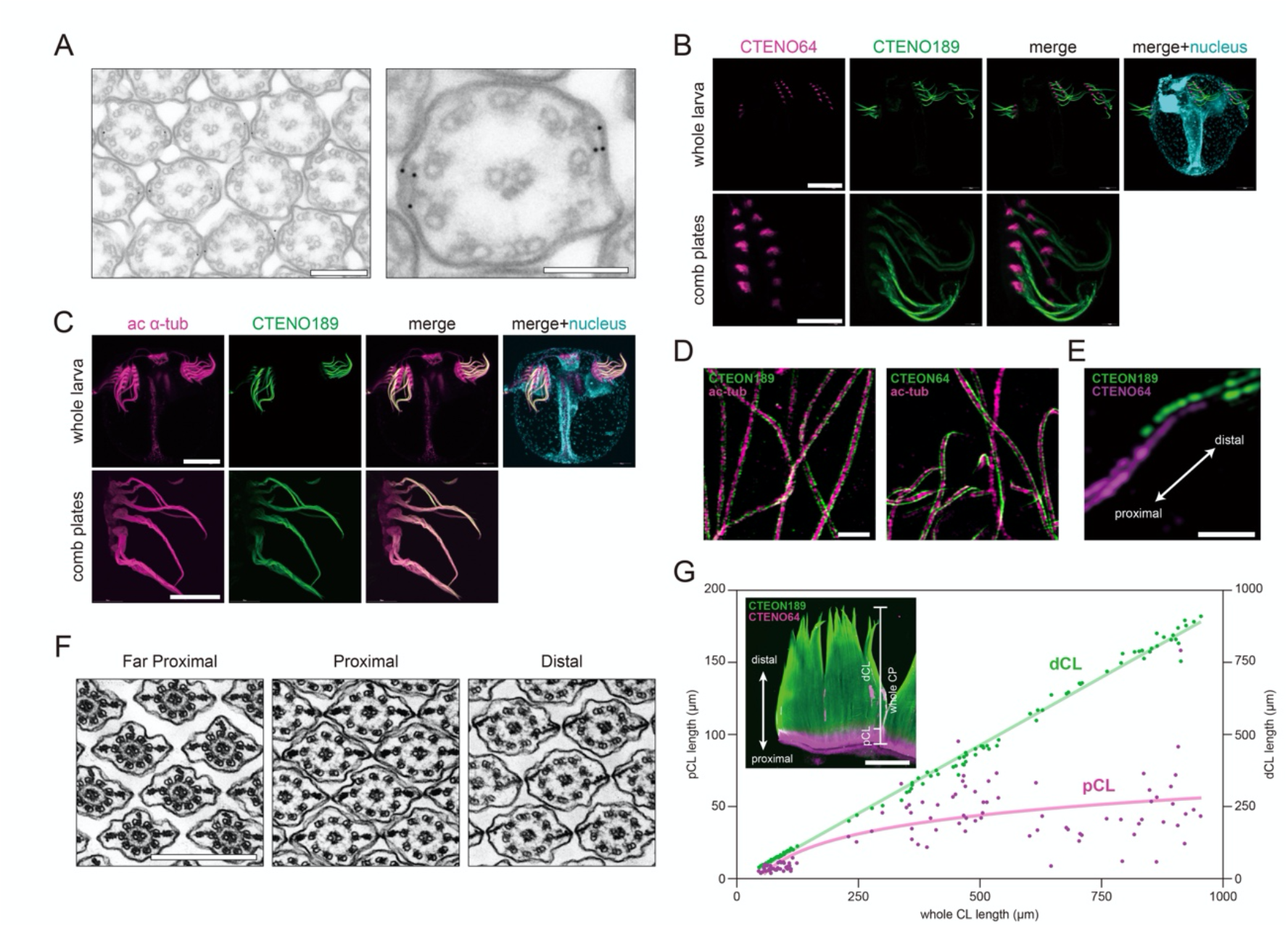
CTENO189 is a CL protein localized at the distal region of comb plates. (A) Left, post-embedding immunogold labeling with an anti-CTENO189 antibody in the distal part of an adult comb plate. Bar, 200 nm. Right, a magnified image. Bar, 100 nm. (B) Immunolocalization of CTENO189 in a whole larva (36 hpf). Double-staining with anti-acetylated α-tubulin antibody (ac α-tub, magenta), anti-CTENO189 antibody (green) and DAPI for nuclei (cyan). Top, whole larva. Bar, 100 μm. Bottom, comb plates. Bar, 50 μm. (C) Double-staining of a whole larva with anti-CTENO64 antibody (magenta), anti-CTENO189 antibody (green) and DAPI for nuclei (cyan). Top, whole larva. Bar, 100 μm. Bottom, comb plates. Bar, 50 μm. (D) Localization of CTENO189 and CTENO64 in a fragmented cilium from an adult comb plate using a Lattice SIM. The axoneme and lateral rows of CLs are clearly distinguished. Left, double-staining of CTENO189 (green) and acetylated α-tubulin (ac-tub, magenta); right, double-staining of CTENO64 (green) and acetylated a-tubulin (ac-tub, magenta). Each protein was localized along two rows at the lateral end of a single cilium. Bar, 500 nm. (E) Double-staining of CTENO189 (green) and CTENO64 (magenta). Two rows of CLs in a cilium are shown. At the boundary of the proximal and distal regions, the two proteins localized completely separately. The arrow indicates the distal and proximal directions in the comb plate. Bar, 1 μm. (F) Sequential images of TEM cross-sections in the proximal CP region. Images at proximal (left), middle proximal (middle), and distal (right) regions of CP are shown. Note that the morphology of CLs and the interciliary space are different between images. Additional CL at doublet 1 microtubule is observed in some cilia of far proximal image. Bar, 500 nm. (G) Growth of proximal and distal CLs. The lengths of proximal and distal CL were measured from double-stained fluorescent images with anti-CTENO64 and anti-CTENO189 antibodies (inset). Data from hatched cydippid larvae (29-56.5 hpf) (triangle), developed larvae with lobes and tentacles (30-47 days, square) and adults (circles) were plotted against the whole CP length. Bar, 200 μm.

To obtain an ultrastructural basis for the segregation between proximal and distal CLs, we prepared serial sections of a CP and examined the arrangement of cilia using transmission electron microscopy (TEM) (Figures 1F and S3). In addition to the difference in morphology between the proximal and distal CLs,^3^ we observed a transition in the ciliary array around the proximal region. From the ciliary rootlets and basal bodies located near the cell surface, multicilia protruded from the polster cell (Figure S3). In the vicinity of the cell surface, the cilia lacked CLs and were arranged in a certain space between them (Figure 1F). CLs in the far proximal region of a CP are found at doublet 1, as well as doublets 3 and 8 (Figure 1F, left). The proximal CL appeared as the diameter of the axonemes increased, and the space between the cilia became confined (Figure 1F, middle). The interciliary space expanded distally (Figure 1F, right). These observations demonstrate that CPs have a distinct compartmentalization in CL morphology and in the interciliary space between the proximal and distal regions of a CP, representing a “two-story building” structure.

We measured the length of the proximal or distal CL stained with anti-CTENO64 or anti-CTENO189 antibodies, respectively (Figure S4). The proximal CL region extended in the early stage of CP growth but remained constant (43.1±17.7 μm, n = 34) after the CP length reached up to ~500 μm (Figure 1G). Further growth of the CP accompanied extension of the distal region of the CP (Figure 1G). Formation of the proximal CL was also observed during regeneration of the excised part of the comb row. After a part of a comb plate row was excised, an interplate ciliary groove (ICG) and a mass of polster cells began to regenerate in 3 days to a week, as reported for *M. leidyi*.^11^ Anti-CTENO64 antibody recognized the proximal CL in the center of polster cells, where the CTENO189-positive distal CL was not observed in the lateral region of CP but was extended from the proximal CL in the middle (Figure S5).

### CTENO189 is required for CL formation in the distal region

CTENO64 was clearly localized in the proximal CL, but its knockdown caused the loss of CL up to only 16% of the comb plate.^3^ To determine the role of CTENO189 in the formation and motility of CPs, we knocked down its gene expression in *B. mikado* larvae using translation blocking morpholinos (MO). We used two MOs (MO1 and MO2) targeted to the 5’-UTRs of *CTENO189* mRNA. We microinjected MOs into fertilized eggs and cultured the embryos for up to 30 hours postfertilization (hpf). Western blotting and immunofluorescence microscopy showed that CTENO189 was completely lost in morphant larvae (Figures 2A and S6A). No significant changes were observed in the morphology of larval organs, including the apical organs, pharynx, ciliated grooves, and CPs (Figure 2A). MO-based knockdown of CTENO64 did not affect the expression or localization of CTENO189 in the larvae (Figure 2B).

**Figure 2.**
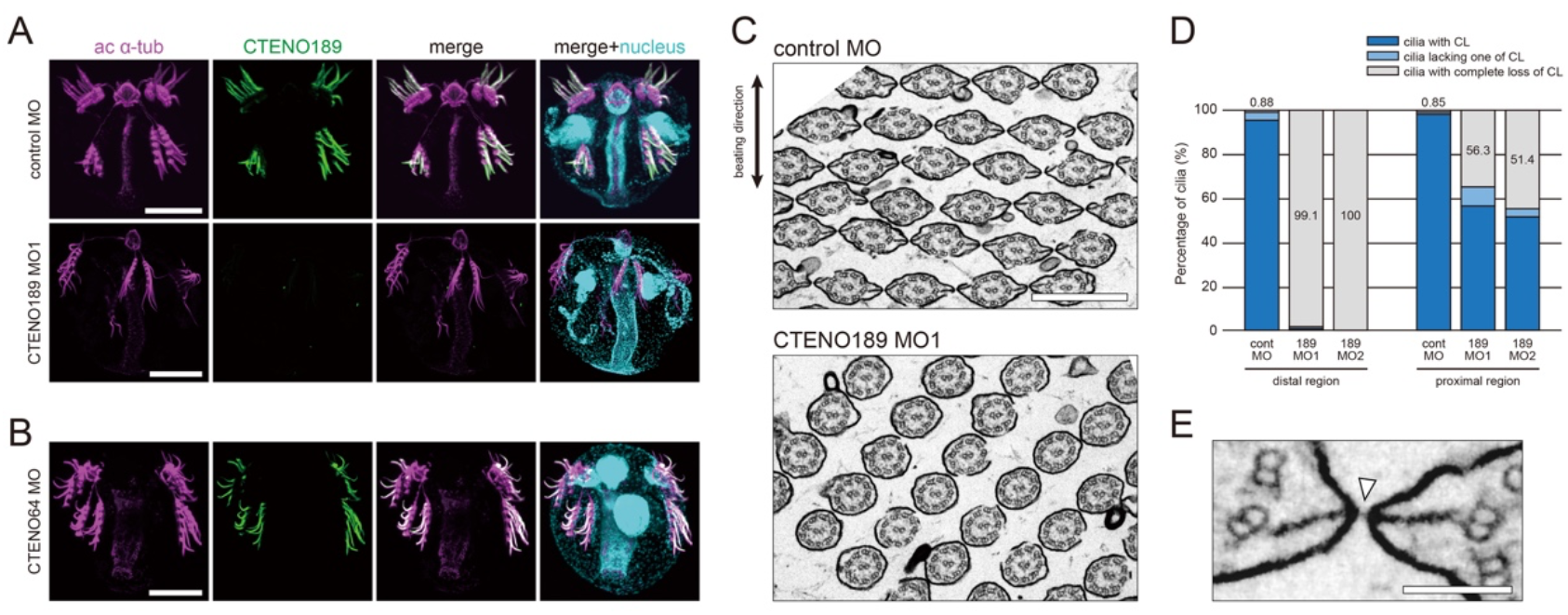
Knockdown of CTENO189 causes loss of the distal CL. (A) Immunofluorescence analysis of CTENO189 between control (control MO) and CTENO189 morphant (CTENO189 MO1), showing the loss of CTENO189 in the distal CL. The larvae were stained with antibodies against acetylated α-tubulin (ac α-tub, magenta) and CTENO189 (green) and with DAPI for nuclei (cyan). Bars, 100 μm. (B) Double-staining of CTENO64 morphant (CTENO64 MO) with antibodies against acetylated α-tubulin (ac α-tub, magenta) and CTENO189 (green) and with DAPI for nuclei (cyan), showing normal formation of distal CL. Bar, 100 μm. (C) TEM images of the distal CP region in the control (upper: control MO) and CTENO189 (lower: CTENO189 MO1) morphant. The arrow on the left indicates the beating direction of the comb plate. Bar, 500 nm. (D) Comparison of the percentage of cilia at the proximal and distal CP regions in the control (contMO) and CTENO189 (189MO1 and MO2) morphants. Dark blue, light blue, or grey represents cilia with CL, cilia lacking one of the CLs, or cilia completely lacking CL, respectively. Data from contMO (n = 791, 10 comb plates), 189MO1 (n = 851, 8 comb plates), and 189MO2 (n = 821, 12 comb plates), pCL (proximal CL): contMO (n = 2817, 10 comb plates), 189MO1 (n = 1005, 5 comb plates), and 189MO2 (n = 1340, 7 comb plates). (E) A magnified image of the CL connection between the cilia in the control larvae. The white arrowhead shows the interciliary bridge that connects the two CLs from adjacent cilia. Bar, 100 nm.

We found that knockdown of the *CTENO189* gene resulted in no significant change in the length of larval CPs (Figure S6B) or the number of cilia per CP area between the control and morphant larvae (Figure S6C). To investigate the effect of CTENO189 knockdown on the ultrastructure of CP, we compared cross sections of CPs between control and morphant larvae using TEM. Although no significant difference was found in the structures of axonemes, nearly all cilia in the distal region of the morphant CP were devoid of CLs (Figures 2C and 2D). More than half of the cilia retained complete CLs in the proximal region of the morphant larval CPs (Figure 2D).

To examine the effect of CL loss on the ciliary array in the distal region, we compared the distribution of CP cilia. Adjacent cilia are linked through interciliary bridges, which extend from CLs to the extracellular region (Figure 2E). In morphant CPs, the interciliary bridge was lost and the distance between cilia in the CP beating plane was widened (Figure 2C). To quantify the arrangement of cilia, we marked the center of the cilia with a dot in the cross-sectioned TEM images of control and morphant larval CP. In the control, cilia were arranged in a regular hexagonal array, whereas the arrangement became disordered with nonconstant interciliary distances in the morphant (Figure 3A). We measured the angle and distance between adjacent cilia in the horizontal and diagonal directions (Figure S7A), and plotted them in polar coordinates (Figure 7B). In the control larvae, two horizontally adjacent cilia in the CP were clustered approximately 350 nm apart, whereas six diagonally adjacent cilia were spaced 250 nm apart and clustered in four diagonal locations at 50°, 130°, −50°, and −130° (Figure 3B; contMO).

**Figure 3.**
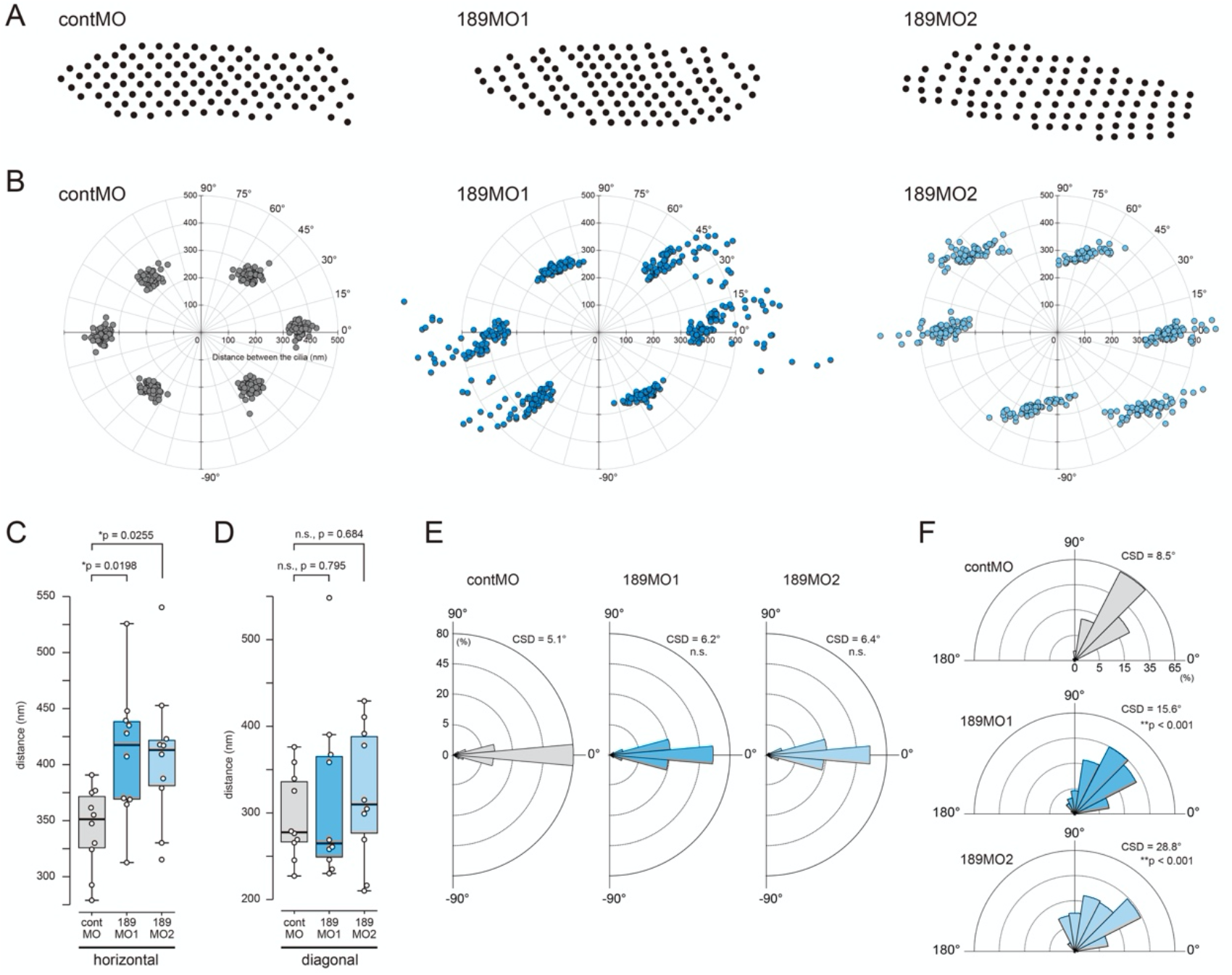
Loss of the distal CL in CTENO189 morphants disrupts hexagonal lattice arrangement of CP cilia. (A) Ciliary arrangement in a CP distal region in the control (contMO) and CTENO189 (189MO1 and MO2) morphants. Dots represent the centers of cilia plotted from TEM cross sections. A typical pattern from ten TEM images is shown for each sample. (B) The distance and the angle between all adjacent cilia in (A) were plotted on polar coordinates. (C) The average distance between the cilia in the horizontal directions was measured in the control and CTENO189 morphants. The average distance was significantly increased in the CP of morphant larvae (*p < 0.05, Dunnett’s multiple-comparison test). (D) The average distance between the cilia in the diagonal directions shows no significant (n.s.) difference between the control and CTENO189 morphants. (E) Circular histograms of the angle between two adjacent cilia in the horizontal direction, showing no significant (n.s.) difference between control and CTENO189 morphants. (F) Circular histograms of the angle between two adjacent cilia in the diagonal direction, showing significant difference between the control and CTENO189 morphants (**p < 0.001, Watson’s U2 test). CSD, circular standard deviation in E and F. Data from contMO (grey; n = 783, 10 comb plates), 189MO1 (dark blue; n =928, 10 comb plates), and 189MO2 (light blue; n = 729, 10 comb plates) are shown in C-F.

In morphant larvae, the hexagonal array of CP cilia was disarranged (Figure 3B; 189MO1, 189MO2). The distance between cilia in the horizontal direction increased in the morphant CP, although no significant difference in distance was observed in the diagonal area (Figures 3C and 3D). We evaluated expressed the angular variation between cilia as a circular standard deviation (CSD). The angles between cilia in the morphant were nearly unchanged from those in the control larvae in the horizontal direction (contMO, CSD = 5.1°; MO1, CSD = 6.2°; MO2, CSD = 6.4°) (Figure 3E). However, those in the diagonal direction showed larger variation (189MO1, CSD = 15.6°; 189MO2, CSD = 28.8°) compared to controls (contMO, CSD = 8.5°; p < 0.001) (Figure 3F).

Loss of the proximal CL component, CTENO64, disrupts the ciliary orientations, that is, the axis of two central microtubules, in CP multicilia.^3^ Therefore, we tested whether the loss of CTENO189 affected the orientation of cilia in the distal region of the CP in control and morphant larvae. The results showed that there was no significant difference in cilia orientation between the control and CTENO189 morphants (Figure S7B).

### Loss of distal CL decreases mechanical resistance which induces persistence of asymmetric ciliary wave

To investigate how the loss of CL in the distal region due to CTENO189 knockdown affects the function of CPs, motility was recorded with a high-speed camera of comb plates in control and CTENO189 morphant larvae for comparative analysis. CP beating was similar between the control and CTENO189 morphant larvae in normal seawater. Therefore, considering the possible role of CL in the mechanical reinforcement of giant multicilia, methylcellulose was added to seawater to increase viscous resistance. The asymmetric waveforms of the morphant comb plates were slightly disturbed in 0.2% methylcellulose seawater compared with the control (Figure 4A, Videos S1 and S2). An increase in the concentration of methylcellulose to 0.5% caused the conversion of CP waveforms to symmetry in CTENO189 morphants (Figure 4A, Video S3). The asymmetric index was estimated as the ratio of the maximum and minimum curvatures in the middle of the total length of the CP. The value was found to be significantly smaller and closer to 1 in the morphants than in the control (Figure 4B), indicating that the morphant comb plates beat symmetrically. However, there was no significant difference in ciliary beat frequency between controls and morphants in normal seawater or seawater containing methylcellulose (Figures 4C).

**Figure 4.**
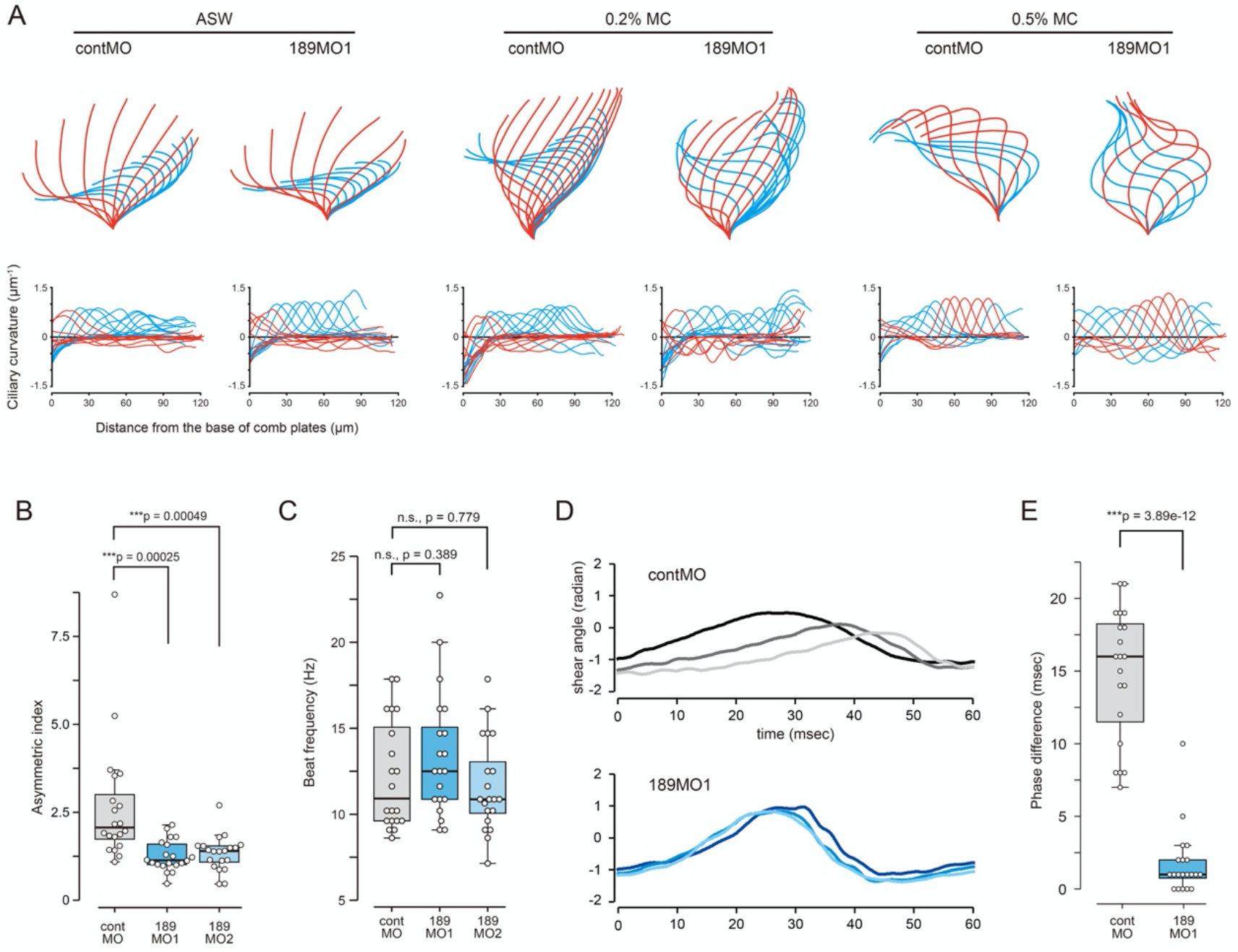
Loss of CL induces symmetrical beating of CPs. (A) Comparison of the CP waveforms between the control (contMO) and CTENO189 (189MO1) morphants. Left, beating waveforms (top) and ciliary curvature along a CP (bottom) in artificial seawater (left), 0.2% methylcellulose seawater (middle) and 0.5% methylcellulose seawater (right). Red and blue represent effective and recovery strokes, respectively. (B) Comparison of the asymmetric index of CPs in 0.5% methylcellulose seawater between control and CTENO189 morphants. The values were measured at the half position of the CP from the base of the CP. Differences between samples were statistically evaluated by Dunnett’s multiple-comparison test (**p < 0.001). (C) CP beat frequency in 0.5% methylcellulose seawater between control and CTENO189 morphants. Differences between samples were statistically evaluated by Dunnett’s multiple-comparison test (n.s., not significant). Data from contMO (gray; n = 20 comb plates, 12 larvae), 189MO1 (dark blue; n = 20 comb plates, 9 larvae), and 189MO2 (light blue; n = 20 comb plates, 15 larvae) are shown in B and C. (D) Changes in the shear angles of three adjacent comb plates in the control (contMO) and CTENO189 (189MO1) morphant. Shear angles at 10 μm from the base of a CP were measured and plotted against time. The different colors of the lines represent the individual CPs. (F) Difference in wave phase between adjacent two CPs were compared between the control (contMO) and CTENO189 (189MO1) morphants. Data are from contMO (n = 20, 9 larvae) and 189MO1 (n = 20, 5 larvae). Differences between samples were statistically evaluated by Welch student’s t test (**p < 0.001).

To investigate the effect of CTENO189 knockdown on the antiplectic metachronal rhythm of CP beating in a comb row, we measured the shear angle of three consecutive CPs in the control and morphant larvae (Figure 4D). The phase differences were maintained in the control larvae in the presence of 0.5% methylcellulose, showing metachronal transmission of beating from one CP to the adjacent. However, it disappeared in the morphant larvae (Figure 4E). These results suggest that the CLs of the distal region are not only important for the maintenance of CP asymmetric waveforms, but also for the establishment of a metachronal rhythm along a comb row.

## DISCUSSION

We previously identified a ctenophore-specific protein, CTENO64,^3^; however, the molecular characterization of the whole CL was only “halfway” achieved, since the localization of this protein is limited to the proximal region of the CL. Here, we identified a novel protein, CTENO189, as another component of CL, from a comprehensive search for CL components by taking advantage of accumulated transcriptome data and information on ciliary and flagellar components. Phylogenomic analyses suggest that ctenophores are a sister group to all other groups of animals, and that these organisms have a mix of shared and unique characteristics.^12^ Primitive species tend to be considered ancestors of complex species. However, elaborate characteristics acquired in various species could be reduced or lost during evolution. The finding of another example of a ctenophore-specific protein, CTENO189, supports the idea that the CP, a multiciliary giant structure, was acquired through convergent evolution in lineage-specific and unique pathways of ctenophores.

CTENO189 was localized in the distal region of the CPs and was completely separated from the proximal CP region where CTENO64 was localized. Although CLs extend from doublet 3 and 8 to link to the adjacent cilia, TEM images showed morphological differences between proximal and distal regions.^3^ The knockdown of CTENO64 caused only partial loss of the proximal CL, and had no effect on the localization of CTENO189,^3^ whereas that of CTENO189 remarkably inhibited the formation of the distal CL, but not the formation of the proximal CL. This suggests that CL formation does not occur continuously along the CP but occurs independently in the proximal and distal regions, which constructs a two-story building structure. The mechanism by which the two-story frame of CL is formed needs to be solved in light of the mechanism for axoneme formation through intraflagellar transport.^13–15^

Cilia in the proximal CP region are tightly arrayed with less interciliary space than that in the distal region, and their orientations are disturbed by the knockdown of the proximal CL component CTENO64.^3^ A basal structure is observed in the growth process of larval CP and the regeneration process of CP in adults in *M. leidyi*,^11,16^ as well as in the formation of macrocilia, another compound cilia in the mouth of Beroid.^17^ This part is composed of multicilia with dynein arms,^3,11,18^ suggesting that the proximal CL is essential not only as a simple base to determine the ciliary orientation but also as a motile part to establish the orientation for CP beating.^3^ This is supported by the fact that ciliary orientation in epithelia is determined by mechanical feedback from ciliary hydrodynamics.^19,20^

The cilia in the distal CP region are arranged with a larger interciliary space. Motility analysis of morphant larvae demonstrated that CTENO189 is important for the hexagonal arrangement of cilia in the distal CP to maintain an asymmetric waveform. CPs receive ten times more inertial resistance than viscous resistance. In fact, the CPs are thought to move under high Re, estimated at 10-300.^21–23^ Because CTENO189 morphants show larger interciliary distances in the horizontal direction without changing the angles between cilia, it is likely that the horizontal connections between cilia by the distal CL is deeply involved in the maintenance and propagation of asymmetric waves, and that the loss of this connection weakens the mechanical reinforcement against hydrodynamic drag in a high Re world. To overcome such a large drag, ctenophores are thought to have convergently acquired the distal CL during the process of enlarging compound cilia.

From a mathematical model where inertial effects occur, asymmetric ciliary waveforms play an important role in maintaining antiplectic metachronal waves.^24,25^ Antiplectic metachronal waves are thought suitable for fast delivery of low-viscosity liquids in a larger Re environment because the distance between the cilia becomes longer during effective stroke.^26–29^ The mechanism of the metachronal coordination of CP beating has been controversial.^30–32^ The ideas are mainly divided into three: simple electric transmission of membrane potentials,^30^ neuronal transmission,^31^ and mechanical transmission.^2,5,32,33^ A modern model gave a plausible interpretation that ICG cilia drive hydromechanical interactions between CPs, which are transmitted to the center base of CP and laterally spread within a CP.^2^ Ctenophores other than lobate species do not possess ICGs, instead hydromechanical coupling between adjacent CP is thought to play a role in metachronal coordination between CPs.^32^ Our present study showed that the knockdown of CTENO189 results in the loss of both distal CL and metachronal waves without affecting ICG. This strongly suggests that hydrodynamic coupling between and within CPs mediated by the distal CL, and not by ICGs, is a primary mechanism for metachronal waves. The hydromechanical force from the adjacent CP is thought to be laterally transmitted within the CP. CTENO189 or the distal CL should provide an elastic interaction between cilia and drive lateral transmission. Intriguingly, proximal CP and ICG commonly possess morphologically similar CLs that are composed of CTENO64. This continuity is key for the flow of mechanical force from the ICG to the base and lateral directions within the CP.

In conclusion, we demonstrated a two-story building that both structurally and functionally supports the multiciliary movement of a CP. Proximal CLs are robust structures that play key roles in the orientation and propagation of planar movement of the CP in the proximal CP region, whereas distal CLs support the flexibility of an enormous number of cilia and contribute a great deal to overcoming mechanical fluid constraints to maintain asymmetric movement and metachronal wave propagation in a giant CP. The presence of multiple ctenophore-specific CL proteins demonstrates that unique and sophisticated molecular architectures of CPs are the result of the independent evolution of ctenophores, which were selected to locomote their bodies by bundling cilia to develop a giant paddle.

## CONTACT FOR REAGENT AND RESOURCE SHARING

Further information and requests for resources and reagents should be directed to and fulfilled by Kazuo Inaba.

## MATERIALS AND METHODS

### MATERIALS AND RESOURCES

Goat anti-Mouse IgG (H+L) Secondary Antibody, HRP (Thermo Fisher Scientific, Cat#62-6520; RRID: AB_2533947); Goat anti-Rabbit IgG (H+L) Highly Cross-Adsorbed Secondary Antibody, Alexa Fluor 546 (Thermo Fisher Scientific, Cat#A-11035; RRID: AB_2534093); Goat anti-Mouse IgG (H+L) Cross-Adsorbed Secondary Antibody, Alexa Fluor 488 (Thermo Fisher Scientific, Cat#A-11001; RRID: AB_2534069); Goat anti-Rabbit IgG (H+L) Cross-Adsorbed Secondary Antibody, Alexa Fluor 488 (Thermo Fisher Scientific, Cat#A-11008; RRID: AB_1431650; Goat anti-Mouse IgG (H+L) Highly Cross-Adsorbed Secondary Antibody, Alexa Fluor 546 (Thermo Fisher Scientific, Cat#A-11030; RRID: AB_2534089); Anti-α-Tubulin Antibody, clone DM1A (Sigma-Aldrich, Cat#05-829; RRID:AB_310035); Acetyl-α-Tubulin (Lys40) (D20G3) XP Rabbit mAb (Cell Signaling Technology, Cat#5335; RRID: AB_10544694); Goat Anti-Mouse IgG (H+L) Antibody, 5 nm Gold Conjugated (BBI Solutions, Cat#EMGMHL5/0.25; RRID: AB_1769168); Mouse anti-CTENO64 antibody (Jokura et al.^3^); BL21(DE3) Competent E. coli (New England Biolabs, Cat# C2527); Enhanced Chemiluminescence kit, ECL Prime (GE Healthcare, RPN2232); Mnemiopsis leidyi protein models (Mnemiopsis Genome Project Portal, Moreland et al.^34^, https://research.nhgri.nih.gov/mnemiopsis/); Neurobase (http://neurobase.rc.ufl.edu/Pleurobrachia); Mnemiopsis leidyi single cell transcriptome data (Sebé-Pedrós et al.^10^, https://www.nature.com/articles/s41559-018-0575-6#additional-information; Supplementary table 4); Mouse: BALB/cAnNCrlCrlj (Charles River Laboratories, JAX: 000651); Primers for CTENO189, 5’-GCGCGGATCCTTTGCTGACAAAAATGAGAT-3’(forward) and 5’-GCGCGAATTCTGAGAAATTGCGGGTATCTT-3’ (reverse); CTENO189-MO1 (Gene Tools, 5’-AGCTCATGGTGGCTGAACTCAAATT-3’); CTENO189-MO2 (Gene Tools, 5’-GTGGCTGAACTCAAATTATTTGTAT-3’); CTENO64-MO (Gene Tools, 5’-ATAGTCAGAGCTTTTCGCACTTCGT-3’; Jokura et al.^3^); Control-MO (Gene Tools, CCTCTTACCTCAGTTACAA TTTATA-3’, Gene Tools; Jokura et al.^3^); pET-28a(+) (Merck Cat# 69864); Software and Algorithms, Scaffold 4; Proteome Software (https://www.proteomesoftware.com/products/scaffold-4); FV10-ASW (Olympus, https://www.olympus-lifescience.com/ja/support/downloads/fv10i_vw_license/); ZEN (Zeiss, https://www.zeiss.co.jp/microscopy/products/microscope-software/zen.html); ImageJ (Schneider et al.^35^; https://imagej.nih.gov/); Bohboh (MEDIA LAND Co.,Ltd.; http://shoji-baba.art.coocan.jp/Bohboh_V4.html); Rstudio (The R Foundation, https://www.r-project.org/); Cutwin (3web, http://www.osk.3web.ne.jp/~cutwin/introidx.htm); GENETYX14 (Genetyx, https://www.genetyx.co.jp/).

### EXPERIMENTAL MODEL AND SUBJECT DETAILS

Adults of the ctenophore *B. mikado* were collected by ladling a dipper from a pier or by snorkeling at Oura Bay (Shimoda), Nanao Bay (Nanao), Tabira Bay (Hirado), and Aburatsubo Bay (Misaki). Adults were maintained in a circulating 80-L aquarium with a constant supply of filtered natural seawater under a light/dark cycle. The temperature of the seawater in the aquarium was maintained at 22°C using a cooler and a heater. They were fed with 20-50 ml of brine shrimp at 75-100 individuals/ml per day. The amount of feed varied according to the number of individuals in the aquarium. To induce sexual maturation, frozen copepods were provided to the adults five times a day for two weeks. Two mature adults were transferred to a 1 L beaker and left to stand in the dark for 5 h, after which light stimulation induced spawning. Fertilized eggs were collected from the bottom of the beaker using a pipette, transferred to a petri dish, and placed in an incubator set at 20°C until use.

#### MS analysis using nanoLC-MS/MS and screening for the candidates of CL proteins

Adult comb plates were cut from their bases using a tungsten needle. Whole proteins of the isolated adult comb plates were separated by SDS-PAGE and stained with CBB. The polyacrylamide gel was cut into 10 small pieces of descending molecular weights and subjected to in-gel digestion with trypsin. Peptide masses were analyzed using nanoLC-MS/MS at the Support Center for Advanced Medical Sciences, Tokushima University. Proteins were identified with Mascot software (version 2.5) using the transcriptome assembly sequence database of *B. mikado*.^3^ The results were analyzed using Scaffold 4 (Proteome Software, OR, USA). All row data for proteomic analysis were deposited in jPOST^36^ with the accession numbers JPST001482 and PXD031689. To screen CL candidates, MS-identified proteins were first selected by normalized exponentially modified protein abundance index (emPAI)^9^ with values greater than 50, using Scaffold 4. These proteins were then subjected to a BLASTP search against the NCBI database and further screened by exclusion of ciliary and flagellar proteins, including dynein-related, radial-spoke-related, ciliary and flagellar associated (CFAP), and coiled-coil domaincontaining (CCDC) proteins. The proteins were then subjected to a local BLAST search against the *M. leidyi* protein model (version 2.2) (http://research.nhgri.nih.gov/mnemiopsis). Single-cell transcriptome data from adult *M. leidyi*^10^ were used for further screening by cell clusters (https://www.nature.com/articles/s41559-018-0575-6#additional-information; Supplementary table 4, *M. leidyi* cell clusters enriched gene lists), in which C48 and C50 clusters are grouped into “comb cells”.^37^ In total, 21 proteins were finally selected.

#### Sequence analysis

A local BLAST search against the databases for ctenophore species was performed using the transcriptomes hosted on Neurobase (https://neurobase.rc.ufl.edu/pleurobrachia/download), which contains data from the ctenophores *Beroe abyssicola, Bolinopsis ashleyi, Bolinopsis infundibulum, Coeloplana astericola, Dryodora glandiformis, Euplokamis dunlapae, Mnemiopsis leidyi, Pleurobrachia bachei, Pleurobrachia pileus, Pukia falcata,* and *Vallicula* multiformis. Domain searches were performed using SMART software (http://smart.embl-heidelberg.de/).

#### Production of antibodies against CTENO189

The open reading frame of the CTENO189 cDNA was amplified by PCR. The primers used were: 5’-GCGCGGATCCTTTGCTGACAAAAATGAGAT-3’ (forward) and 5’-GCGCGAATTCTGAGAAATTGCGGGTATCTT-3’ (reverse). The amplified cDNA and pET 28a (+) were cleaved with *Eco*RI and *Bam*HI, purified using an S-400 spin column (GE Healthcare, IL, USA), ligated, and electroporated into *E. coli* BL21. After checking for transformation, bacteria from a single colony were inoculated into LB medium with 50 μg/ml kanamycin and incubated at 37°C. Protein expression was induced using 1 mM isopropyl-1-thio-β-d-galactopyranoside during the logarithmic growth phase. The bacteria were harvested by centrifugation, and the recombinant CTENO189 was purified according to a previously described protocol.^37^ After dialysis against PBS, recombinant CTENO189 was emulsified with Freund’s complete adjuvant and immunized into female BALB/c mice by three subcutaneous injections at intervals of 10 days. Blood tests were performed before antiserum collection. Rabbit anti-CTENO189 polyclonal antibody was prepared by Biologica Corporation, Nagoya, Japan.

#### Western blot analysis

The proteins from the isolated adult comb plates or whole larvae were separated by SDS-PAGE and transferred to polyvinylidene difluoride membranes. Membranes were treated with 7.5% skim milk in PBS containing 0.05% Tween 20 (PBST). Blots were incubated with anti-CTENO189 antibody or anti-CTENO64 antibody^3^ at a 1:1000 dilution, or anti-α-tubulin antibody (05-829, Millipore, Darmstadt, Germany) at a 1:10,000 dilution for 1 h at room temperature (RT). After washing with PBST, the blots were incubated with an anti-mouse HRP-conjugated secondary antibody at a 1:10,000 dilution for 30 min at RT. After washing with PBST three times, the blots were developed using an enhanced chemiluminescence kit (ECL Prime, GE Healthcare, IL, USA).

#### Immunofluorescence microscopy

Immunofluorescence microscopy was performed using a protocol previously reported,^3^ with some modifications. The whole body of the hatched larvae was used as a sample. For larvae, a comb row was cut out with a tungsten needle. For adults, a single comb plate was cut and used. As for the regenerating comb plates, the comb plates that were regenerated 3 days to 1 week after removing a part of a comb row of adults were used as samples according to the method of Tamm^1.11^ These samples were fixed with cold methanol containing 50 mM EGTA at −30°C for 30 min. The samples were washed with excess PBST and incubated in blocking solution (10% goat serum and 1% BSA in PBS) for 1 h at RT, followed by incubation with anti-CTENO189 antibody and anti-acetylated-α-tubulin antibody (T6793, Sigma-Aldrich, MD, USA) or anti-CTENO64 antibody^3^ at 1:1000 dilution in the blocking solution overnight at 4°C. After washing with PBST at RT, the samples were incubated with Alexa 488 goat anti-mouse IgG and Alexa 546 goat anti-rabbit IgG or Alexa 488 goat anti-rabbit IgG and Alexa 546 goat anti-mouse IgG at a 1:1000 dilution in blocking solution for 6 h at RT. The samples were washed with PBST at RT and incubated with 10 μM DAPI in PBS. Images were obtained using an Fv10 confocal microscope (Olympus).

#### Measurement of CP length

The length of the whole CP and those of the proximal and distal CL regions were measured for three to five positions per adult CP or five CPs from each larva, respectively, from confocal immunofluorescent images stained with both anti-CTENO64 and anti-CTENO189 antibodies. The three longest curve paths along the ciliary line of a CP were traced using the line tool of ImageJ, with a line width of 20 or 1 for adult and larval CP, respectively. The fluorescence intensity along the line was plotted against the distance for both colors. The boundary between the areas of the two different fluorescent colors was defined as the intersection point between the two curves (Figure S4). The lengths of the proximal and distal CL were plotted against the whole CP length (Figure 1G).

#### Super-resolution microscopy

For super-resolution imaging of individual cilia, the isolated adult CPs were passed through a 25 gauge needle several times for fragmentation and disintegration, immobilized on a glass slide, and processed for immunostaining. by using the procedure described above. Super-resolution images were obtained by structured illumination microscopy (lattice SIM) using an Elyra 7 (ZEISS, Germany).

#### Immunogold labeling

Adult comb plates were cut from their bases using a tungsten needle. For post-embedding immunogold labelling, the isolated adult comb plates were attached to a glass slide precoated with 1 mg/ml poly-l-lysine. After 5 min at RT, the comb plates were fixed with 4% paraformaldehyde (PFA), 0.02% glutaraldehyde, 0.4 M sucrose, 0.1 M cacodylate buffer (pH 7.4). Samples were washed several times with 0.1 M cacodylate buffer (pH 7.4) and post-fixed with 1% OsO_4_, 1.5% sodium ferrocyanide, 0.1 M cacodylate buffer (pH 7.4) for 30 min on ice. After washing with 0.1 M sodium cacodylate (pH 7.4), the samples were dehydrated through a graded ethanol series and embedded in LR-white at 55°C for 48 h. The block was thin-sectioned with an average thickness of 70 nm and deposited on a nickel grid (300 mesh) supported by formvar films. The sections were pretreated for etching with a saturated aqueous solution of sodium metaperiodate for 1 min, followed by five washes in distilled water. They were then sequentially treated with 1% BSA in PBS for 1 h, incubated with primary anti-CNENO189 antibody (1:1000 dilution) at 4°C overnight, washed four times with PBS, and incubated with antimouse secondary antibody conjugated with 5 nm gold (1:50 dilution, BBI, UK) for 1 h. After washing with PBS, sections were stained with 5% aqueous uranyl acetate for 5 min and observed under a 1200EX electron microscope (JEOL, Tokyo, Japan) at 80 kV.

#### Microinjection of morpholino antisense oligonucleotide (MO)

CTENO189 MO1 (5’-AGCTCATGGTGGCTGAACTCAAATT-3’) and CTENO189 MO2 (5’-GTGGCTGAACTCAAATTATTTGTAT-3’) were designed to cover the 5’-UTR, including the start codon of *B. mikado CTENO189* mRNA (Gene Tools, Philomath, OR, USA). Each MO was dissolved in distilled water, dispensed in small quantities, and stored at 4°C until further use. An aliquot of MO was heated at 65°C for 10 min before use to ensure complete dissolution. Microinjections were performed with pressure using a previously reported protocol.^3^ The MO was dissolved in 400 mM KCl and 20% glycerol to 1 mM and injected into fertilized eggs with vitelline membranes before cleavage. The injection volume was controlled by measuring the diameter of each droplet. The final concentration of MO in the egg was ~3.5 μM. The eggs were kept in filtered seawater at 20°C for further development.

#### Electron microscopy for larval comb plates

The control and morphant larvae were fixed with 2.5% glutaraldehyde in 0.45 M sucrose, 0.1 M sodium cacodylate (pH 7.4), and washed with 0.1 M sodium cacodylate (pH 7.4). Fixed larvae were embedded in a pre-warmed 1.5% low-melt agarose gel. After coagulating agarose, the samples were trimmed and post-fixed with 1% OsO_4_ at 4°C for 1 h. After dehydration in a graded ethanol series, samples were embedded in Agar Low Viscosity Resin (Agar Scientific, Stansted, UK) through propylene oxide and thin-sectioned with an average thickness of 70 nm. The sections were stained with uranyl acetate and lead citrate and observed under a 1200EX electron microscope (JEOL, Tokyo, Japan) at 80 kV.

#### Analysis of ciliary orientation

TEM images of the cross sections of comb plates between control and morphant larvae were used to analyze ciliary orientations. The angles between the plane of the central pair and ciliary beating plane were measured. The data were compared using circular histograms as described previously.^3^

#### Analysis of ciliary alignment

TEM images of comb plate cross-sections were used to analyze the ciliary alignment in control or morphant larvae. The cilia adjacent to the comb plates were arranged in a hexagonal array. The plane of two central-pair microtubules of the cilium was used as a reference. The adjacent cilia located parallel to the central pair plane were designated as horizontal, and those located toward the beating direction as diagonal (Figure S4A). The distance (l) was measured between each center of adjacent cilia, where a cilium had adjacent cilia in six directions (two horizontal and four diagonal directions) in the hexagonal array. The angle (d) between the cilia was measured in the direction perpendicular to the beating direction, defined as 0°, using the ImageJ software.

#### Analysis of ciliary beating

The injected larvae at 30 hpf were placed in a 35-mm plastic glass-based dish at 20°C. Motility of the comb plates was recorded using an inverted microscope (IX71, Olympus, Tokyo, Japan) equipped with a high-speed camera (500 frames per second; HAS-U2, DITECT, Tokyo, Japan). The waveforms were traced and analyzed using the Bohboh software (Bohboh Soft, Tokyo, Japan), as described previously.^3^ The asymmetry of ciliary movement was expressed as the ratio of maximum positive and negative curvatures. This ratio should be 1 when the comb plate shows a completely symmetric waveform, and becomes greater than 1 when it shows an asymmetric waveform. This ratio is referred to as the “asymmetric index”.^3^

### QUANTIFICATION AND STATISTICAL ANALYSIS

Statistical analysis is described in Figures (p values and group sizes) and their legends (statistical test uses, definition of n).

## ACKNOWLEDGEMENTS

We thank the staff of Notojima Aquarium, Hakkeijima Sea Paradise, Kujukushima Aquarium, Misaki Marine Biological Station (University of Tokyo), International Coastal Research Center, and the Atmosphere and Ocean Research Institute (University of Tokyo) for their cooperation in collecting and rearing the ctenophores. This work was supported in part by a Grant-in-Aid for Scientific Research (B) No. 22370023 and for Challenging Research (Pioneering) No.20K20583 from the Japan Society for the Promotion of Science, Japan (JSPS) to K.I., Grant-in-Aid for the JSPS Research Fellow 18J10941 to K.J., and by a Grant-in-Aid for Innovative Areas No. 22370023, 25291069, and 15H01201, from the Ministry of Education, Culture, Sports, Science and Technology, Japan. We would like to thank Editage (www.editage.com) for English language editing.

## AUTHOR CONTRIBUTIONS

K.I. designed and supervised the research; K.J. established the rearing system for ctenophores; K.J., Y.S., K.S., and K.I. performed the research; K.J. and K.I. wrote the manuscript; and all authors agreed to the final version of the manuscript.

## DECLARATION OF INTERESTS

The authors declare no competing interests.

## DATA AND CODE AVAILABILITY

The transcriptome dataset of *B. mikado* supporting the current study is available in the DDBJ Sequenced Read Archive under accession code DRA008845. The accession number for the CTENO189 sequence reported in this study is GenBank LC681832. Data for proteomic analysis are available at jPOST^36^ under accession numbers JPST001482 and PXD031689.

## Notes

### Competing Interest Statement

The authors have declared no competing interest.

## REFERENCES

1. Afzelius, B. A. (1961). The fine structure of cilia from ctenophore swimming-plates. J. Biophys. Biochem. Cytol. 9, 383–394.

2. Tamm, S. L. (2014). Cilia and the life of ctenophores. Invertebr. Biol. 133, 1–46.

3. Jokura, K., Shibata, D., Yamaguchi, K., Shiba, K., Makino, Y., Shigenobu, S., and Inaba, K. (2019). CTENO64 is required for coordinated paddling of ciliary comb plate in ctenophores. Curr. Biol. 29, 3510–3516.e4.

4. Dentler, W. L. (1981). Microtubule-membrane interactions in cilia and flagella. Int. Rev. Cytol. 72, 1–47.

5. Tamm, S. L. (1984). Mechanical synchronization of ciliary beating within comb plates of ctenophores. J. Exp. Biol. 113, 401–408.

6. Welch, V. L., Vigneron, J. P., and Parker, A. R. (2005). The cause of colouration in the ctenophore *Beroe cucumis*. Curr. Biol. 15, R985–R986.

7. Horridge, G. A. (1964). The giant mitochondria of ctenophore comb plates. Q. J. Cell Sci. s3-105, 301–310.

8. Tamm, S. L., and Tamm, S. (1981). Ciliary reversal without rotation of axonemal structures in ctenophore comb plates. J. Cell Biol. 89, 495–509.

9. Ishihama, Y., Oda, Y., Tabata, T., Sato, T., Nagasu, T., Rappsilber, J., and Mann, M. (2005). Exponentially modified protein abundance index (emPAI) for estimation of absolute protein amount in proteomics by the number of sequenced peptides per protein. Mol. Cell. Proteomics 4, 1265–1272.

10. Sebé-Pedrós, A., Chomsky, E., Pang, K., Lara-Astiaso, D., Gaiti, F., Mukamel, Z., Amit, I., Hejnol, A., Degnan, B. M., and Tanay, A. (2018). Early metazoan cell type diversity and the evolution of multicellular gene regulation. Nat. Ecol. Evol. 2, 1176–1188.

11. Tamm, S. L. (2012). Regeneration of ciliary comb plates in the ctenophore *Mnemiopsis leidyi*. I. Morphology. J. Morphol. 273, 109–120.

12. Dunn, C. W., Leys, S. P., and Haddock, S. H. (2015). The hidden biology of sponges and ctenophores. Trends Ecol. Evol. 30, 282–291.

13. Rosenbaum, J. L., and Witman, G. B. (2002). Intraflagellar transport. Nat. Rev. Mol. Cell Biol. 3, 813–825.

14. Ishikawa, H., and Marshall, W. F. (2017). Intraflagellar transport and ciliary dynamics. Cold Spring Harb. Perspect. Biol. 9, a021998.

15. Morthorst, S. K., Christensen, S. T., and Pedersen, L. B. (2018). Regulation of ciliary membrane protein trafficking and signalling by kinesin motor proteins. FEBS Journal 285, 4535–4564.

16. Tamm, S. L. (2012). Patterns of comb row development in young and adult stages of the ctenophores *Mnemiopsis leidyi* and *Pleurobrachia pileus*. J. Morphol. 273, 1050–1063.

17. Tamm, S. L., and Tamm, S. L. (1988). Development of macrociliary cells in *Beroe*. II. Formation of macrocilia. J. Cell Sci. 89, 81–95.

18. Jokura, K., and Inaba, K. (2020). Structural diversity and distribution of cilia in the apical sense organ of the ctenophore *Bolinopsis mikado*. Cytoskeleton (Hoboken) 77, 442–455.

19. Mitchell, B., Jacobs, R., Li, J., Chien, S., and Kintner, C. (2007). A positive feedback mechanism governs the polarity and motion of motile cilia. Nature 447, 97–101.

20. Mizuno, K., Shiba, K., Yaguchi, J., Shibata, D., Yaguchi, S., Prulière, G., Chenevert, J., and Inaba, K. (2017). Calaxin establishes basal body orientation and coordinates movement of monocilia in sea urchin embryos. Sci. Rep. 7, 10751.

21. Blake, J. R., and Sleigh, M. A. (1974). Mechanics of ciliary locomotion. Biol. Rev. Camb. Philos. Soc. 49, 85–125.

22. Matsumoto, G. I. (1991). Swimming movements of ctenophores, and the mechanics of propulsion by ctene rows. Hydrobiologia 216-217, 319–325.

23. Barlow, D., Sleigh, M. A., and White, R. J. (1993). Water flows around the comb plates of the ctenophore *Pleurobrachia* plotted by computer: A model system for studying propulsion by antiplectic metachronism. J. Exp. Biol. 177, 113–128.

24. Knight-Jones, E. W. (1954). Relatio ns between metachronism and the direction of ciliary beat in Metazoa. J. Cell Sci. s3–95, 503–521.

25. Sleigh, M. A. (1963). Movements and co-ordination of the ciliary comb plates of the ctenophores *Beroe* and *Pleurobrachia*. Nature 199, 620–621.

26. Dong, X., Lum, G. Z., Hu, W., Zhang, R., Ren, Z., Onck, P. R., and Sitti, M. (2020). Bioinspired cilia arrays with programmable nonreciprocal motion and metachronal coordination. Sci. Adv. 6, eabc9323.

27. Hussong, J., Breugem, W. P., and Westerweel, J. (2011). A continuum model for flow induced by metachronal coordination between beating cilia. J. Fluid Mech. 684, 137–162.

28. Goebel, W. H., Colin, S. P., Coatello, J. H., Bemmell, B. J., and Sutherland, K. R. (2020). Scaling of ctenes and consequences for swimming performance in the ctenophore *Pleurobrachia bachei*. Invertebr. Biol. 139, e12297.

29. Colin, S. P., Costello, J. H., Sutherland, K. R., Gemmell, B. J., Dabiri, J. O., and Du Clos, K. T. (2020). The role of suction thrust in the metachronal paddles of swimming invertebrates. Sci. Rep. 10, 17790.

30. Horridge, G. A. (1965). Intracellular action potentials associated with the beating of the cilia in ctenophore comb plate cells. Nature 205, 602.

31. Sleigh, M. A. (1972). Features of ciliary movement of the ctenophores *Beroe, Pleurobrachia* and *Cestus*. In: Essays in Hydrobiology, Clark, R. B. and Wootton, R. (eds.), Exeter University Press, Exeter, pp. 119–136.

32. Tamm, S. L. (1973). Mechanisms of ciliary coordination in ctenophores. J. Exp. Biol. 59, 231–245.

33. Moss, A. G., and Tamm, S. L. (1986). Electrophysiological control of ciliary motor responses in the ctenophore *Pleurobrachia*. J. Comp. Physiol. A 158, 311–330.

34. Moreland, R. T., Nguyen, A. D., Ryan, J. F., Schnitzler, C. E., Koch, B. J., Siewert, K., Wolfsberg, T. G., and Baxevanis, A. D. (2014). A customized Web portal for the genome of the ctenophore *Mnemiopsis leidyi*. BMC Genom. 15, 316.

35. Schneider, C. A., Rasband, W. S., and Eliceiri, K. W. (2012). NIH Image to ImageJ: 25 years of image analysis. Nat. Methods 9, 671–675.

36. Okuda, S., Watanabe, Y., Moriya, Y., Kawano, S., Yamamoto, T., Matsumoto, M., Takami, T., Kobayashi, D., Araki, N., and Yoshizawa, A. C., Tabata, T., Sugiyama, N., Goto, S., and Ishihama, Y. (2017). jPOSTrepo: An international standard data repository for proteomes. Nucleic Acids Res. 45, D1107–D1111.

37. Padma, P., Satouh, Y., Wakabayashi, K., Hozumi, A., Ushimaru, Y., Kamiya, R., and Inaba, K. (2003). Identification of a novel leucine-rich repeat protein as a component of flagellar radial spoke in the Ascidian *Ciona intestinalis*. Mol. Biol. Cell 14, 774–785.

