## Supplementary material for "“Two-story building” of a ctenophore comb plate provides structural and functional integrity for motility of giant multicilia": Figure S1, Figure S2, Figure S3, Figure S4, Figure S5, Figure S6, Figure S7, Table S1, Table S2, Table S3

SUPPLEMENTARY MATERIALS

SUPPLEMENTARY FIGURES

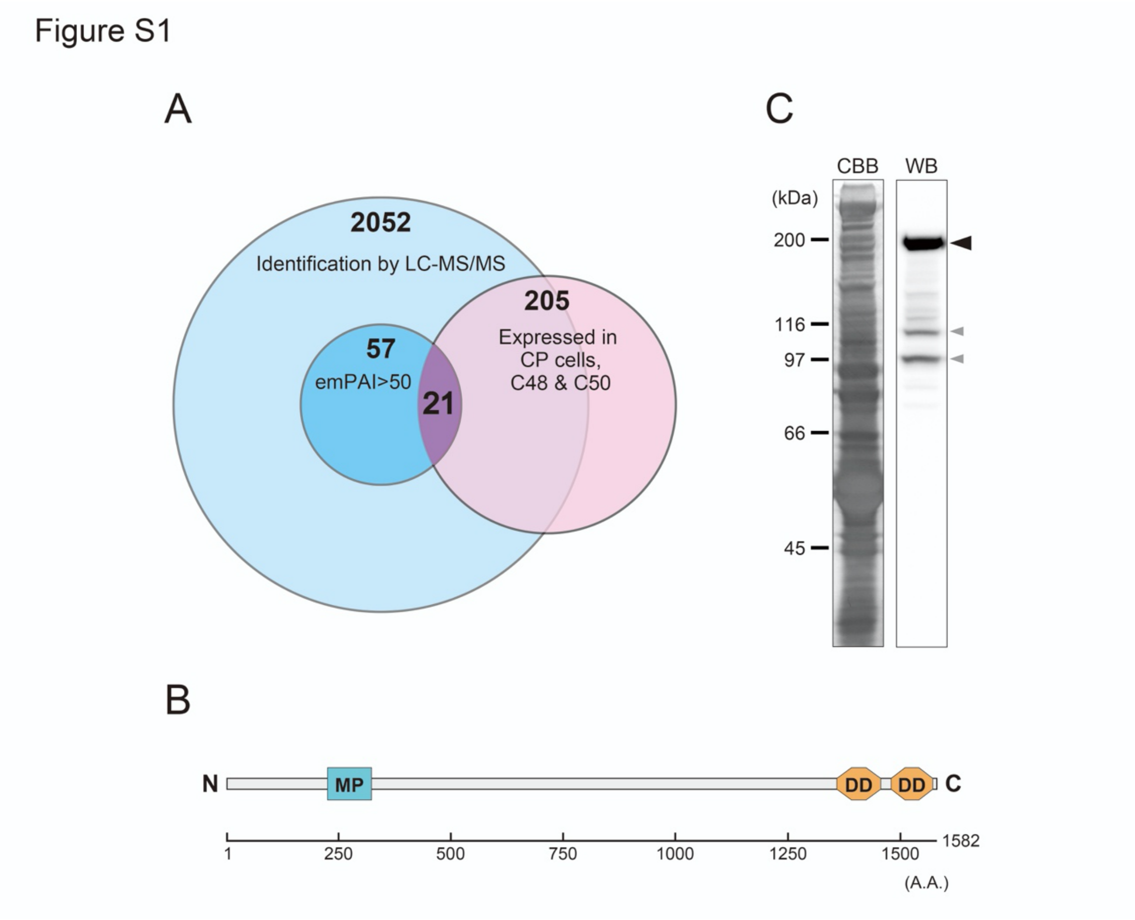


**Figure S1. Identification and characterization of CTENO189.**

(A) Strategy for screening of CL proteins in *B. mikado*. The candidates for CL components were screened for high abundance in CPs and exclusive gene expression in CP cells.

(B) The domain structure of CTENO189 predicted by SMART. The protein is a 1582 amino acid (A.A.) protein with two death domains (DD, orange) from A.A.1359 to 1457 (E-value, 3.87) and A.A.1480 to 1573 (E-value, 1.20e-7), and a zinc-dependent metalloprotease domain (MP, blue) from A.A.224 to 322 (E-value, 8.00e-11). N and C represents the N- and C-termini, respectively.

(C) Western blot analysis of whole CP proteins from adult *B. mikado*. The 189-kDa protein was specifically detected (black arrowhead). Gray arrowheads indicate degradation products of the 189-kDa protein. CBB, Coomassie brilliant blue staining; WB, western blot.

Figure S2

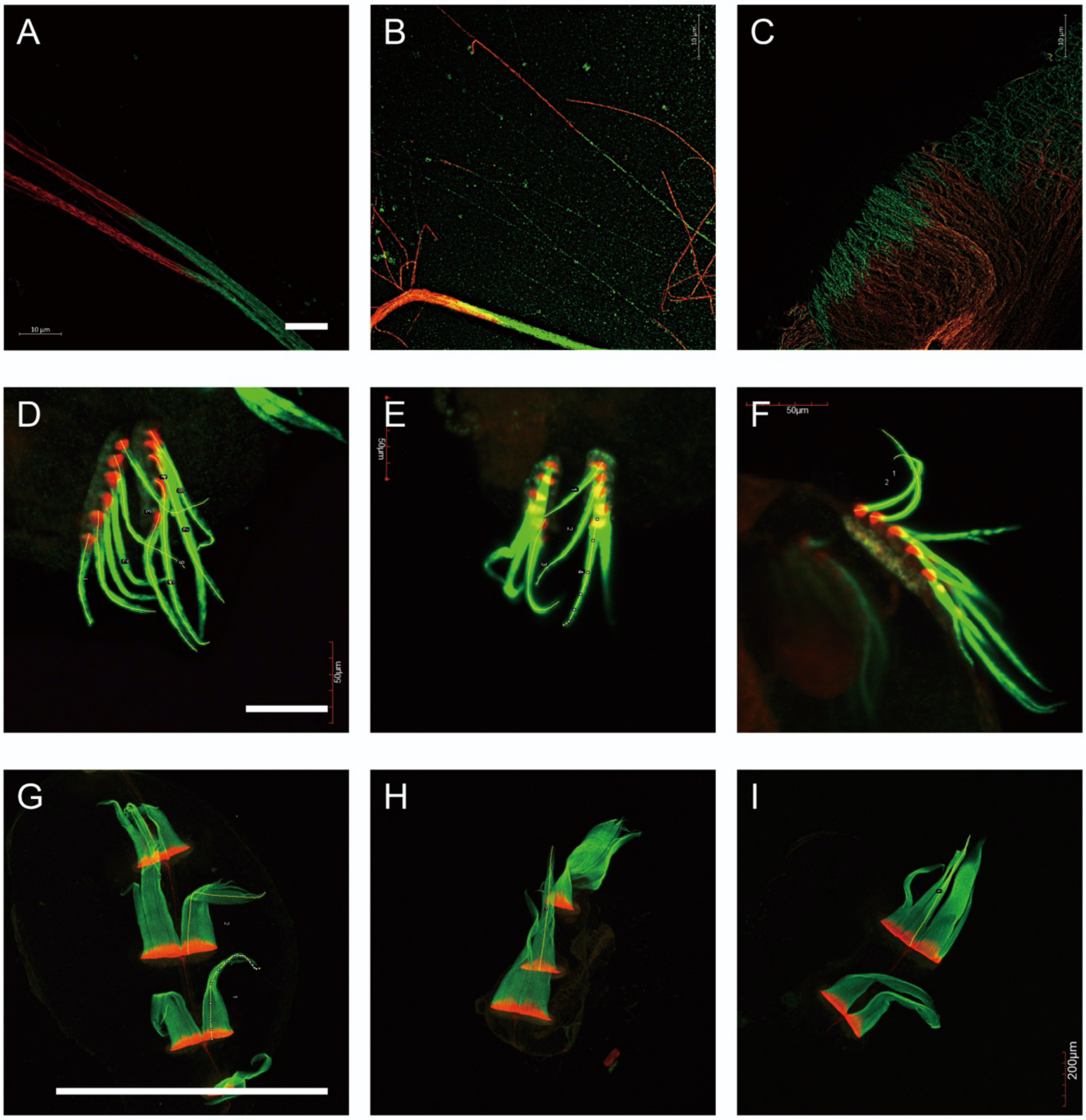


**Figure S2. A gallery for immunofluorescence images of adult and larval CPs.**

Isolated adult and larval CPs were double-stained with anti-CTENO64 (magenta) and anti-CTENO189 (green) antibodies, respectively.

(A–C) Images of adult CPs that were fragmented and disintegrated using a needle. Bar, 10 µm.

(D–F) Images of CPs in hatched cydippid larvae (29-56.5 hpf). Bar, 50 µm.

(G–I) Images of CPs in developed larvae with lobes and tentacles (30-47 days). The apical part of the larvae was excised and processed for immunostaining. Bar, 1 mm.

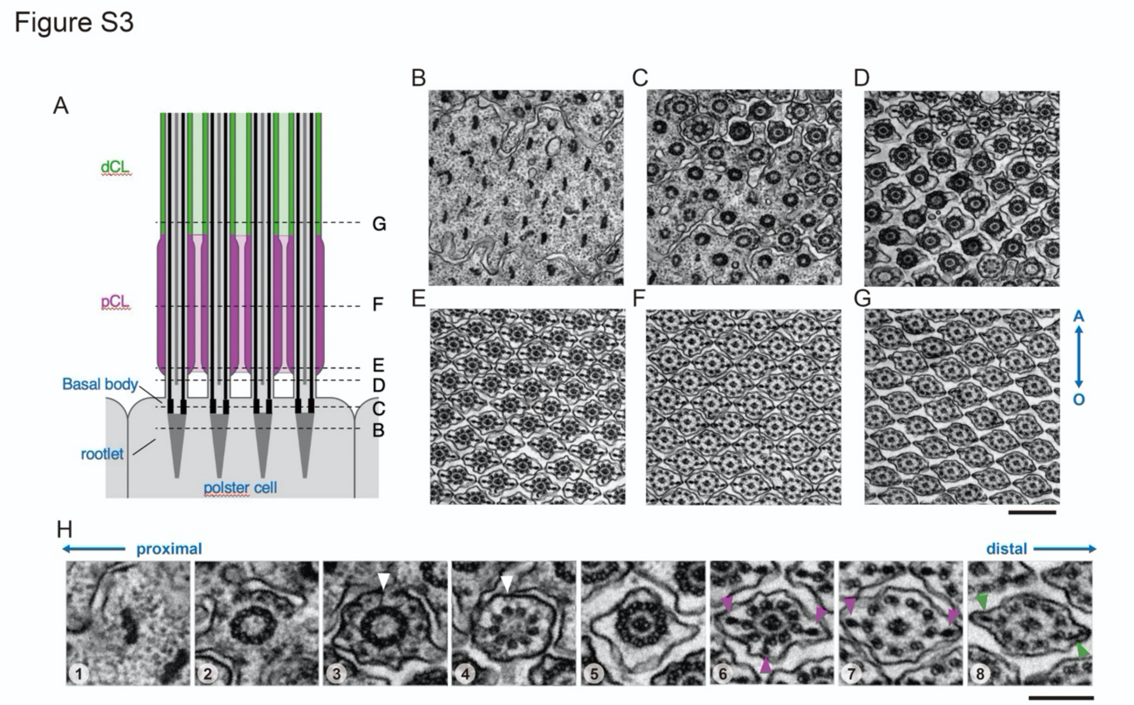


**Figure S3. Ultrastructural transition of a CP from the base to the proximal and distal regions.**

(A) Schematic drawing of the base and proximal region of a CP. Magenta or green represents proximal (pCL) or distal CL (dCL), respectively. Light colors indicate the regions of the ciliary bridge. The positions indicated by dashed lines correspond to the TEM images from (B) to (G).

(B–G) Sequential images of TEM cross sections. The top and bottom of each image show the aboral and oral sides of the CP, respectively. Bar, 500 nm.

(H) Sequential images of the cilium at several positions in the CP. 1, rootlet; 2, basal body; 3, basal body with transition fibers (arrowhead); 4, transition zone with Y-links (arrowhead) attached to doublet microtubules; 5, most distal part of the axoneme with 9+2 structure without CLs; 6, beginning of proximal CP region with CLs (arrowhead; CL from doublet microtubule 1 is often observed); 7, distal CP region with thickened CLs (arrowhead); 8, beginning of proximal CP region with thinner and longer CLs (arrowhead). Note that the extracellular space between cilia is altered among the regions.

Figure S4

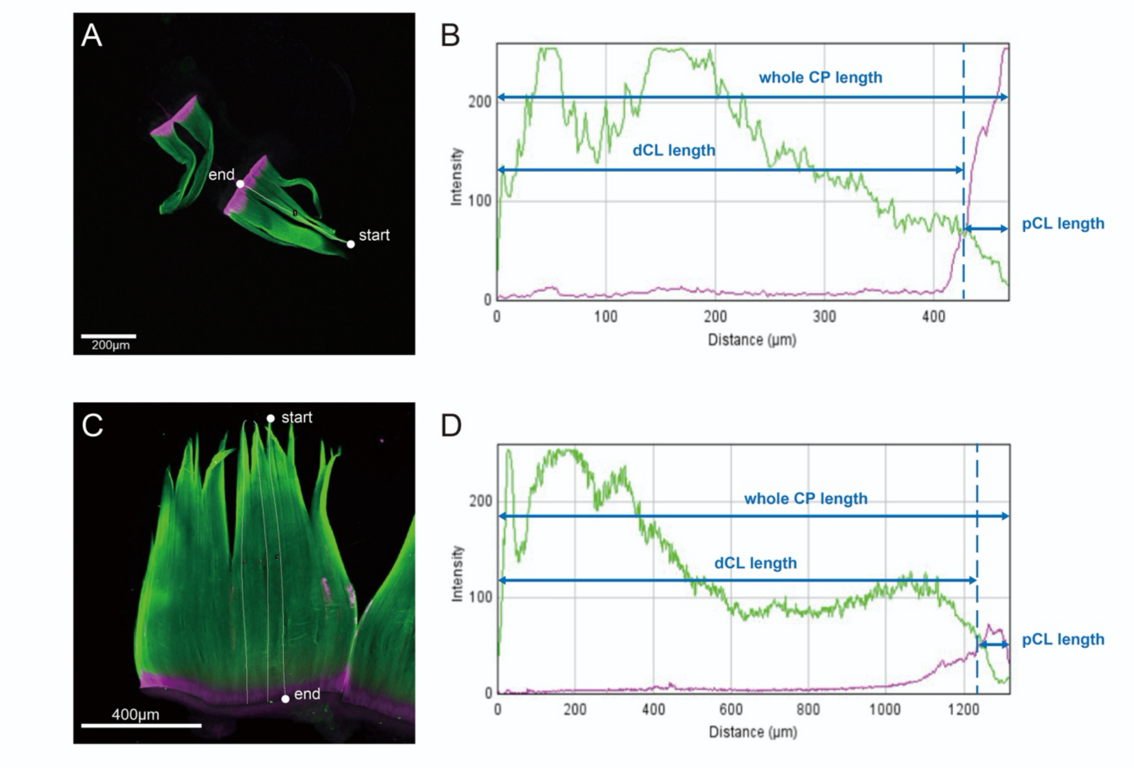


**Figure S4. Measurement of CP and CL lengths.**

Double-stained images of larval (A) and adult (C) CPs are shown with the line of the curve path used for measurement by ImageJ. The starting and ending points for the measurement are shown as dots. (B, D) Definition of the length of the entire CP, distal CL (dCL), and pCL (proximal CL). The fluorescence intensity along the curve path was plotted against the distance from the starting point (tip of the CP). Bars, 200 μm (A) and 400 μm (B).

Figure S5

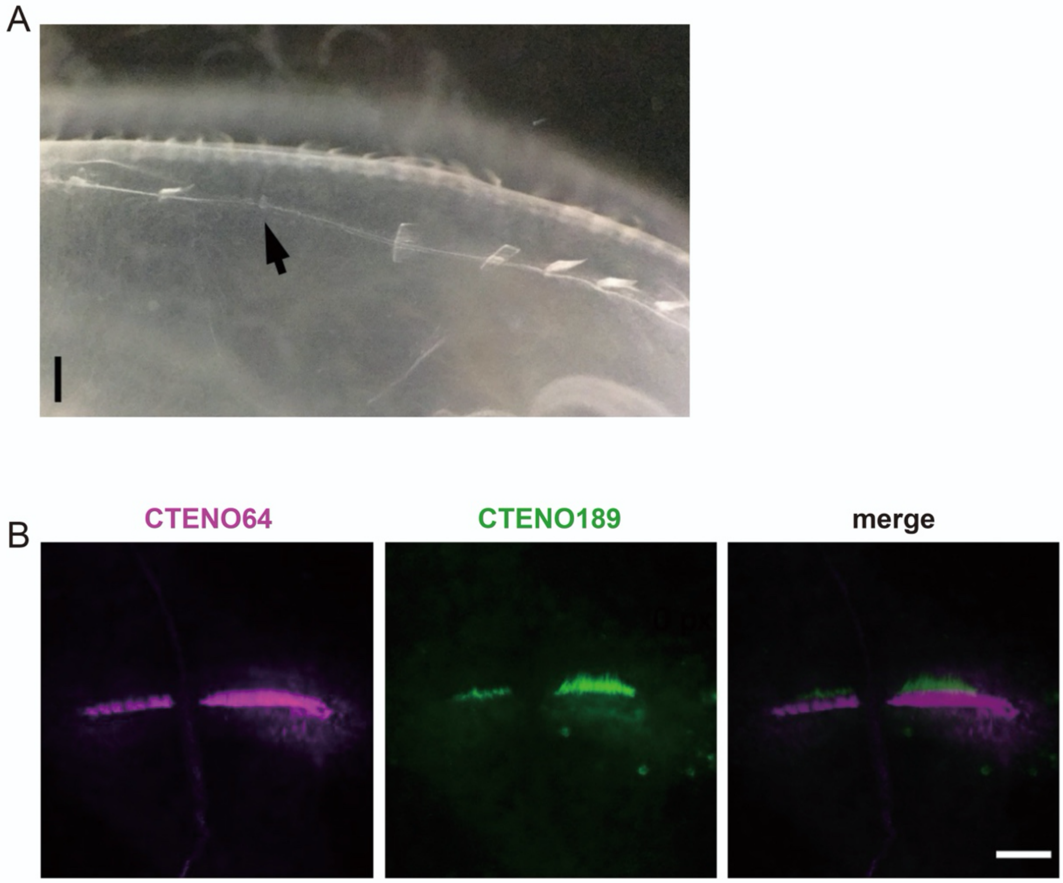


**Figure S5. Immunofluorescence image of CLs in a regenerating CP.**

(A) Regeneration of a comb plate. A newly formed CP regenerated from an excised part of a comb row is shown by an arrow. Bar, 2 mm.

(B) Magnified image of double-stained CP. Proximal CL develops noticeably at this stage. Localization of proximal (CTENO64) and distal (CTENO189) CLs are segregated from an early stage of CP formation. Bar, 50 µm.

Figure S6

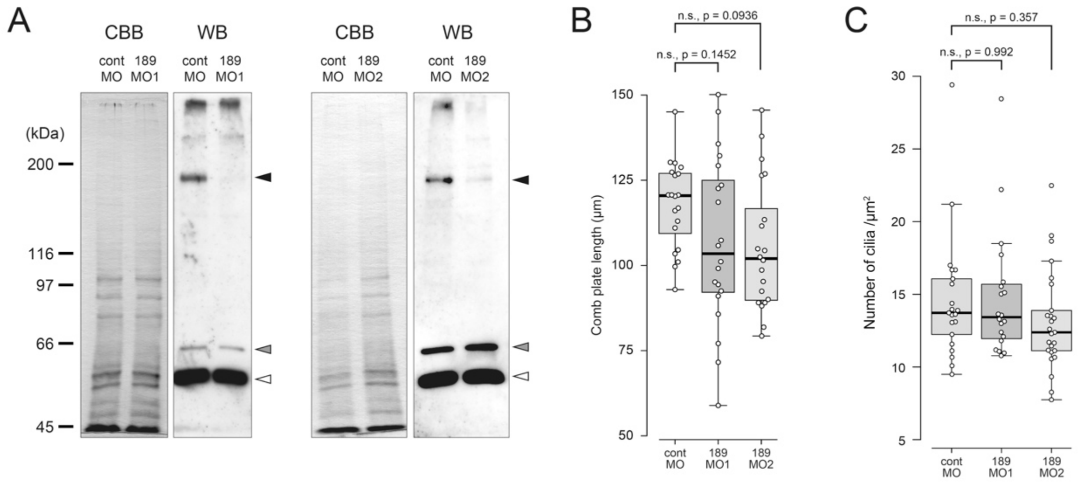


**Figure S6. Analysis of CPs in CTENO189 morphants.**

(A) Immunoblot analysis of whole proteins in the morphant larvae by anti-CTENO189 (black arrowheads), anti-CTENO64 (gray arrowheads), and anti-α-tubulin antibodies (white arrowheads). Left, morphants from CTENO189 morpholino 1 (189MO1); right, morphants from CTENO189 morpholino 2 (189MO2). Proteins from 16 or 8 morphant larvae at 30 hpf were used for 189MO1 or 189MO2, respectively. CBB, Coomassie brilliant blue staining; WB, western blot.

(B) Lengths of the comb plates between control (contMO) and CTENO189 morphant larvae (189MO1 and 189MO2) show no difference. Data were from contMO (20 CPs), 189MO1 (20 CPs), and 189MO2 (20 CPs), and tested by Dunnett's multiple-comparison. n.s., not significant.

(C) Number of cilia per area (1 $\mu m^2$) in a CP between the control (contMO) and CTENO189 morphant larvae (189MO1 and 189MO2) show no difference. Data were from contMO (21 comb plates), 189MO1 (20 comb plates), and 189MO2 (25 comb plates), and tested by Dunnett's multiple-comparison. n.s., not significant.

Figure S7

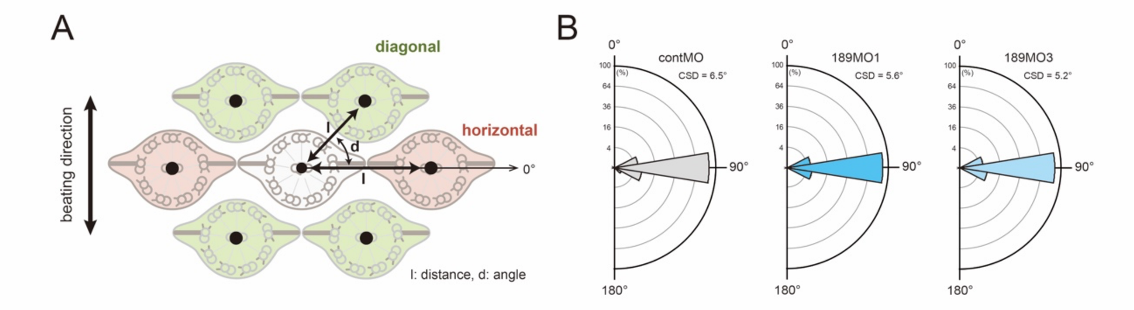


**Figure S7. Ciliary arrangement in CTENO189 morphants.**

(A) Analysis of ciliary array in a CP. A dot was set at the center of two central pair microtubule in each cilium. The distances (l) between adjacent cilia are measured both in the horizontal (red) and diagonal (green) directions. The angle (d) is defined as between horizontal and diagonal lines. The arrow on the left indicates the CP beating direction.

(B) Circular histograms of ciliary orientation in a CP. No significant difference was observed in the orientation of CP cilia between the control larvae and CTENO189 morphants. Data were from contMO (n = 373 from 6 CPs, CSD = 6.5°), 189MO1 (n =318 from 8 CPs, CSD = 5.6°), and 189MO2 (n = 383 from 8 CPs, CSD = 5.2°).

### SUPPLEMENTARY TABLES

**Table S1. Screening of candidates for CL proteins using LC-MS/MS and single-cell transcriptome data.**

| Bm ID | kDa | emPAI | ML homolog | E-value | Identities (%) | Cell types |
|---|---|---|---|---|---|---|
| c55870_g2_i2 | 28 | 771.77 | ML45392a | 9.6E-164 | 97.5 | C28, C48, C49, C50 |
| c52918_g1_i1 | 35 | 424.57 | ML17371a | 0.0 | 99.7 | C48, C50 |
| c46841_g1_i1 | 29 | 395.83 | ML046517a | 0.0 | 96.9 | C48, C49, C50 |
| c57141_g1_i1 | 59 | 243.67 | ML049618a | 0.0 | 97.4 | C48, C50 |
| c57722_g1_i1 | 510 | 222.61 | ML002216a | 0.0 | 98.7 | C27, C28, C48, C49, C50 |
| c50918_g1_i1 | 35 | 192.37 | ML18208a | 0.0 | 99.1 | C48, C49, C50 |
| c50885_g1_i1 | 62 | 163.82 | ML020046a | 0.0 | 94.6 | C26, C27, C28, C48, C49, C50 |
| c57222_g1_i1 | 522 | 147.67 | ML07114a | 0.0 | 97.9 | C27, C48, C50 |
| c57057_g1_i1 | 108 | 140.99 | ML00219a | 0.0 | 96.1 | C48, C50 |
| c55407_g1_i2 | 94 | 137.49 | ML10515a | 0.0 | 95.0 | C28, C48, C50 |
| c54572_g1_i1 | 94 | 107.69 | ML073030a | 0.0 | 96.1 | C31, C48, C49, C50 |
| c55486_g1_i1 | 81 | 107.61 | ML020047a | 0.0 | 97.7 | C48, C49, C50 |
| c47888_g1_i1 | 27 | 102.16 | ML03453a | 6.4E-167 | 93.4 | C31, C48, C49, C50 |
| c50082_g1_i2 | 52 | 85.07 | ML047948a | 0.0 | 98.0 | C48, C49, C50 |
| c57121_g1_i1 | 294 | 77.71 | ML007814a | 0.0 | 94.3 | C48, C49, C50 |
| c56364_g1_i1 | 181 | 70.49 | ML45843a | 0.0 | 93.8 | C48, C49, C50 |
| c56615_g1_i1 | 186 | 67.09 | ML29974a | 0.0 | 95.9 | C48, C50 |
| c51513_g1_i1 | 44 | 65.79 | ML033234a | 0.0 | 97.1 | C48, C49, C50 |
| c49551_g1_i1 | 56 | 60.78 | ML073030a | 7.4E-117 | 91.5 | C31, C48, C49, C50 |
| c54387_g1_i1 | 62 | 52.61 | ML005710a | 0.0 | 97.9 | C27, C48, C49, C50 |
| c48580_g1_i1 | 22 | 51.70 | ML13814a | 6.3E-131 | 99.0 | C48, C50 |

Bm ID: gene ID in the *B. mikado* transcriptome. emPAI, exponentially modified protein abundance index. The ML homolog represents gene ID in *M. leidyi*.

**Table S2. BLASTP homology search of 21 CL candidates.**

| Bm ID | Top Hit-NCBI BLASTP | Species | E-value | Identities (%) |
|---|---|---|---|---|
| c55870_g2_i2 | putative WD repeat-containing protein 65-like | *Apostichopus japonicus* | 9.0E-73 | 54.3 |
| c52918_g1_i1 | PREDICTED: BTB/POZ domain-containing protein 19-like | *Amphimedon queenslandica* | 2.0E-91 | 48.9 |
| c46841_g1_i1 | enkurin-like | *Acropora millepora* | 6.0E-105 | 61.4 |
| c57141_g1_i1 | comb plate cilium protein CTENO64 | *Bolinopsis mikado* | 0.0 | 100.0 |
| c57722_g1_i1 | dynein beta chain, ciliary-like | *Pocillopora damicornis* | 0.0 | 70.3 |
| c50918_g1_i1 | hsp40 heat shock protein 40 | *Phallusia mammillata* | 6.0E-151 | 66.5 |
| c50885_g1_i1 | predicted protein | *Nematostella vectensis* | 2.0E-93 | 39.2 |
| c57222_g1_i1 | dynein heavy chain 5, axonemal-like isoform X1 | *Acanthaster planci* | 0.0 | 63.0 |
| c57057_g1_i1 | coiled-coil domain-containing protein 40-like | *Acropora millepora* | 0.0 | 48.9 |
| c55407_g1_i2 | IQ and AAA domain-containing protein 1-like | *Orbicella faveolata* | 0.0 | 65.2 |
| c54572_g1_i1 | hypothetical protein DRR08_29750 | *Gammaproteobacteria bacterium* | 4.0E-73 | 33.1 |
| c55486_g1_i1 | dynein regulatory complex protein 1 isoform X4 | *Nematostella vectensis* | 0.0 | 51.6 |
| c47888_g1_i1 | PREDICTED: outer dense fiber protein 3-like | *Hydra vulgaris* | 1.0E-64 | 47.5 |
| c50082_g1_i2 | tektin-1 | *Exaiptasia pallida* | 7.0E-70 | 36.2 |
| c57121_g1_i1 | dynein heavy chain 7, axonemal-like isoform X5 | *Acanthaster planci* | 0.0 | 70.0 |
| c56364_g1_i1 | hypothetical protein | *Apibacter sp. HY041* | **2.0E-18** | 31.9 |
| c56615_g1_i1 | cilia- and flagella-associated protein 43-like | *Actinia tenebrosa* | 0.0 | 38.5 |
| c51513_g1_i1 | RIB43A-like with coiled-coils protein 2 | *Nematostella vectensis* | 2.0E-134 | 56.2 |
| c49551_g1_i1 | uncharacterized protein LOC113682194 | *Pocillopora damicornis* | 1.0E-26 | 28.1 |
| c54387_g1_i1 | cilia- and flagella-associated protein 53-like | *Lingula anatina* | 4.0E-140 | 49.5 |
| c48580_g1_i1 | PREDICTED: dynein light chain 1, axonemal | *Dipodomys ordii* | 4.0E-91 | 71.1 |

CTENO189 is shown in red. Bm ID: gene ID in the *B. mikado* transcriptome.

**Table S3 List of orthologous genes for CTENO189 in other ctenophore species.**

| Species | ID | E-value | Identities (%) |
|---|---|---|---|
| *Beroe abyssicola* | Ba_17017_c0_seq2 | 0.0 | 62 |
| *Bolinopsis ashleyi* | comp19144_c0_seq2 | 0.0 | 78 |
| *Bolinopsis infundibulum* | Bi_71906_c1_seq1 | 0.0 | 61 |
| *Coeloplana astericola* | comp18243_c0_seq1 | 3.0E-38 | 64 |
| *Dryodora glandiformis* | sb\|308405\| | 0.0 | 63 |
| *Euplokamis dunlapae* | sb\|10680374\| | 0.0 | 52 |
| *Mnemiopsis leidyi* | Ml_55352_c0_seq26 | 1.0E-118 | 81 |
| *Pleurobrachia bachei* | sb\|9915608\| | 1.0E-148 | 72 |
| *Pleurobrachia pileus* | Pp_45886_c0_seq7 | 0.0 | 48 |
| *Pukia falcata* | comp28930_c0_seq1 | 0.0 | 52 |
| *Vallicula multiformis* | sb\|369518\| | 2.0E-72 | 97 |

A BLAST search against the transcriptome database for several ctenophore species was performed at Neurobase (https://neurobase.rc.ufl.edu/pleurobrachia/download).

**SUPPLEMENTAL MOVIES**

**Video S1.** High-speed video showing propagation of comb plate waveforms in artificial seawater. High-speed video of comb plates of control (left) and CTENO189 morphant (right) larvae beating. The video was recorded at a speed of 0.06×. Bar, 50 µm.

**Video S2.** High-speed video showing propagation of comb plate waveforms in artificial seawater containing 0.2% methylcellulose. High-speed video of comb plates of control (left) and CTENO189 morphant (right) larvae beating. The video was recorded at a speed of 0.06×. Bar, 50 µm.

**Video S3.** High-speed video showing propagation of comb plate waveforms in artificial seawater containing 0.5% methylcellulose. High-speed video of comb plates of control (left) and CTENO189 morphant (right) larvae beating. The video was recorded at a speed of 0.06×. Bar, 50 µm.
